# Constrained neighboring-sarcomere phase topology relates to mean HSO amplitude in living cardiomyocytes

**DOI:** 10.64898/2026.03.13.711515

**Authors:** Seine A. Shintani

**Affiliations:** Department of Biomedical Sciences, College of Life and Health Sciences, Chubu University, Kasugai, Aichi 487-8501, Japan; Center for Mathematical Science and Artificial Intelligence, Chubu University, Kasugai, Aichi 487-8501, Japan; Institute for Advanced Research, Nagoya University, Nagoya, Aichi 464-8601, Japan

**Keywords:** cardiomyocyte, sarcomere, neighboring-sarcomere phase topology, neighboring-sarcomere coordination, weighted synchrony, amplitude-synchrony summary, hyperthermal sarcomeric oscillation

## Abstract

How neighboring sarcomeres redistribute timing while a cardiomyocyte continues to beat, and whether that coordination during warming-induced hyperthermal sarcomeric oscillations (HSOs) is random or structured, remain unresolved. We reanalyzed sarcomere-length recordings from five consecutive sarcomeres in each of seven living neonatal rat cardiomyocytes and represented each valid time point by the four neighboring-pair phase relations that define a 16-state local phase network. During HSOs, the fraction of time with trackable local phase relations increased from 0.298 before warming to 0.956 (paired Wilcoxon *P* = 0.0156), enabling direct analysis of local reconfiguration. Successive local states were almost always connected by Hamming-1 edges, meaning that only one neighboring-pair relation changed at a time (34/35, 97.1%, before warming; 216/230, 93.9%, during HSOs). HSOs also increased occupancy of anti-phase-rich states with three or more anti-phase neighboring pairs (0.254 to 0.509, *P* = 0.0156). These results indicate that HSOs do not reflect unstructured local disorder but a constrained neighboring-sarcomere phase topology.

As a complementary cycle-level analysis within the same HSO window, we then asked how the observed fast amplitude of the valid-sarcomere mean trace relates to local amplitude and synchrony measured from the same valid sarcomeres. For each cycle, *Y*_valid_, the peak-to-peak HSO amplitude of the valid-sarcomere mean trace, was closely approximated by the product of mean local HSO amplitude (*A*) and weighted synchrony (*R_w_*; pooled *r* = 0.992, normalized mean squared error = 0.015, *β*_1_ = 0.948, *β*_0_ ≈ 0). A simple state-derived synchrony factor computed from the local phase patterns showed a modest positive association with *R_w_* (cell-adjusted *β* = 0.197, *P* = 0.0165), providing a bridge between the binary local-state description and the continuous synchrony summary. In blocked cross-validation, the *A × R_w_* summary was markedly more parsimonious than an additive current-state alternative (pooled normalized mean squared error 0.0138 vs 0.1006), whereas simple history terms changed error only marginally. Thus, the main result is a constrained local phase topology during HSOs, and *A × R_w_* serves as a descriptive cycle-level summary of the mean fast signal in the same observed segment, with local amplitude and synchrony as its two components.

## 1 Introduction

Adjacent sarcomeres in a cardiomyocyte are mechanically coupled, yet they need not shorten and relengthen in perfect synchrony. Direct measurements show substantial sarcomere-to-sarcomere heterogeneity in resting and beating cardiomyocytes, and stretch can reduce that heterogeneity by harmonizing strain across the cell (Li et al., 2023; Lookin et al., 2023). In vivo nanoimaging likewise indicates that sarcomeric synchrony and asynchrony are physiologically relevant, influence ventricular pump performance, and remain evident under living myocardial conditions (Kobirumaki-Shimozawa et al., 2021, 2024). Local coordination among neighboring sarcomeres is therefore part of cardiac mechanics itself, not merely a technical complication.

More broadly, cardiac contractility is a multiscale problem in which stochastic molecular events must be converted into ordered cell- and organ-level output. Recent reviews emphasize that this bridge must be understood across scales from single molecules to whole hearts and identify sarcomeres and myofibrils as key intermediate levels in that chain (Garg et al., 2024; Rassier and Månsson, 2025). A quantitative description at this intermediate scale is therefore biologically valuable, because it can connect local contractile heterogeneity to the averaged observables by which beating cells are commonly characterized.

This problem becomes especially sharp in oscillatory contractile states. Partial activation can generate spontaneous tension or length oscillations in striated muscle (Fabiato and Fabiato, 1978; Linke et al., 1993). Under isotonic conditions, shortening and yielding can occur synchronously across sarcomeres (Yasuda et al., 1996), whereas in other settings adjacent yielding clusters or sarcomere “give” can arise and propagate along neighboring elements (Shimamoto et al., 2009; Stehle, 2017). Reviews of cardiomyocyte spontaneous oscillatory contraction (SPOC) have therefore emphasized waves, succession of lengthening phases, and inter-sarcomere coordination as central dynamical features (Kagemoto et al., 2015). What remains unresolved is whether local timing differences among neighboring sarcomeres in living cardiomyocytes reflect unstructured disorder or a constrained mode of reorganization.

Warming provides a useful living-cell context in which that distinction becomes experimentally accessible. Local heat pulses can trigger cardiomyocyte contraction without detectable calcium transients (Oyama et al., 2012), microscopic heating can activate cardiac thin filaments directly (Ishii et al., 2019), and thermal activation shifts thin-filament regulatory equilibria toward activation (Ishii et al., 2020). Cardiac thin-filament activation is also strongly cooperative, involving Ca^2+^, cross-bridge feedback, titin-associated effects, and crosstalk with thick-filament regulatory states (Solís and Solaro, 2021; Park-Holohan et al., 2021). More broadly, length- and strain-dependent activation show that local sarcomere mechanics can feed back onto activation and force generation (de Tombe et al., 2010; Campbell, 2011). In warmed cardiomyocytes, beat-scale rhythm persists while rapid internal oscillations become prominent, creating an experimentally tractable window onto neighboring-sarcomere coordination.

Previous work established warming-induced high-frequency sarcomeric oscillations in living neonatal cardiomyocytes (Shintani et al., 2015), preservation of cycle timing across the HSO state (Shintani et al., 2020; Shintani, 2022), and periodic-chaotic features of that state (Shintani, 2025). What has remained unresolved is both the local spatial organization of neighboring sarcomeres during those oscillations and how that local coordination shapes the oscillatory amplitude seen in the segment-average signal. Here we reanalyzed primary sarcomere-length time series from five consecutive sarcomeres in each of seven warmed cardiomyocytes. Because the post-HSO segment is short, the principal state-space comparisons were restricted to before warming and the HSO segment, whereas the complementary cycle-level analysis was performed within a trimmed HSO window. This reanalysis therefore addresses a mesoscale question: whether neighboring sarcomeres during HSOs traverse local phase space randomly or through a constrained topology, and whether a simple within-segment amplitude-synchrony summary can bridge local coordination and the observed mean fast signal. The reanalysis showed that neighboring sarcomeres do not rephase at random during HSOs. Instead, their trajectories are dominated by minimal single-pair switches within a constrained local phase topology. A simple synchrony factor derived from those local states relates this topology to the continuous synchrony term used in the cycle-level analysis, and mean HSO amplitude is closely approximated by a corresponding amplitude-synchrony summary within the same observed segment.

## 2 Results

### 2.1 Primary sarcomere-length time series reveal a beat-preserving oscillatory regime during HSOs

We first visualized the raw sarcomere-length time series in an exemplar cell to show what changes when the cell enters the HSO state (Figure 1). Before warming, the local 5-sarcomere mean trace showed a comparatively smooth beat envelope and the high-frequency internal component appeared only intermittently. During HSOs, the beat envelope remained visible, but neighboring sarcomeres exhibited sustained rapid internal oscillations superimposed on it. The raw data therefore suggest that the key issue is not whether the mean beat persists, but how neighboring sarcomeres redistribute their relative timing while that beat-scale envelope is maintained.

**Figure 1:**
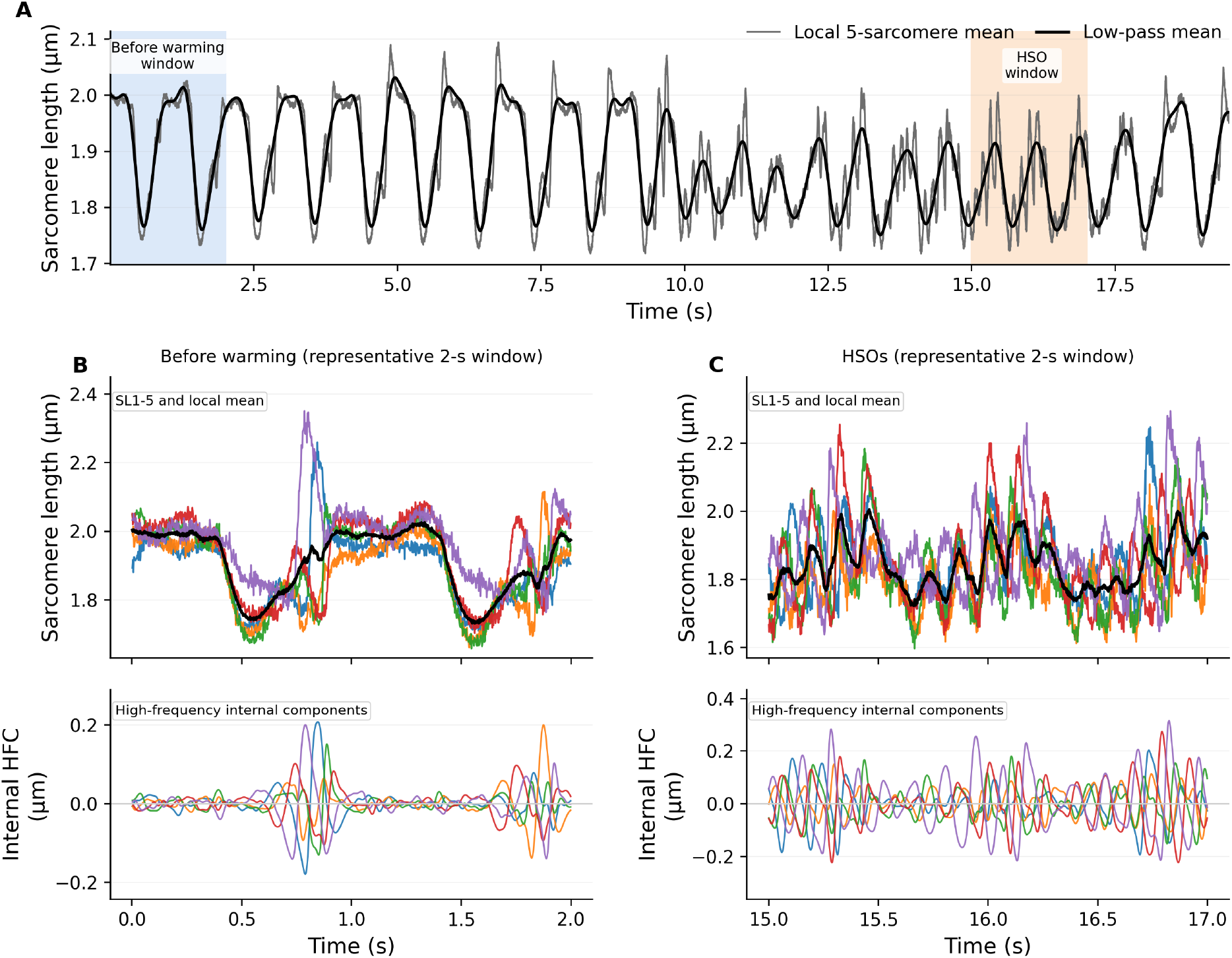
Primary sarcomere-length time-series context before warming and during HSOs in an exemplar cell. **A**, full-trace local 5-sarcomere mean trace for cell 3, with a low-pass mean overlaid. Shaded windows indicate the representative before-warming and HSO segments shown below. **B**, representative before-warming window. Top, sarcomere-length traces of five consecutive sarcomeres and their local mean; bottom, corresponding high-frequency internal components after subtraction of the instantaneous local mean. **C**, representative HSO window shown in the same format.

### 2.2 Local phase relations become trackable through most of the HSO segment

Under the fixed primary gate used for all main-text quantification, the valid fraction for local phase tracking increased from 0.298 before warming to 0.956 during HSOs (paired Wilcoxon *P* = 0.0156; Figure 3A). HSOs therefore provide a regime in which neighboring-pair phase relations can be followed through most of the segment rather than only in brief bursts. A refined visualization gate with longer gap fill was used only to make the representative segment in Figure 2 visually continuous and was evaluated separately in Supplementary Figure S1.

**Figure 2:**
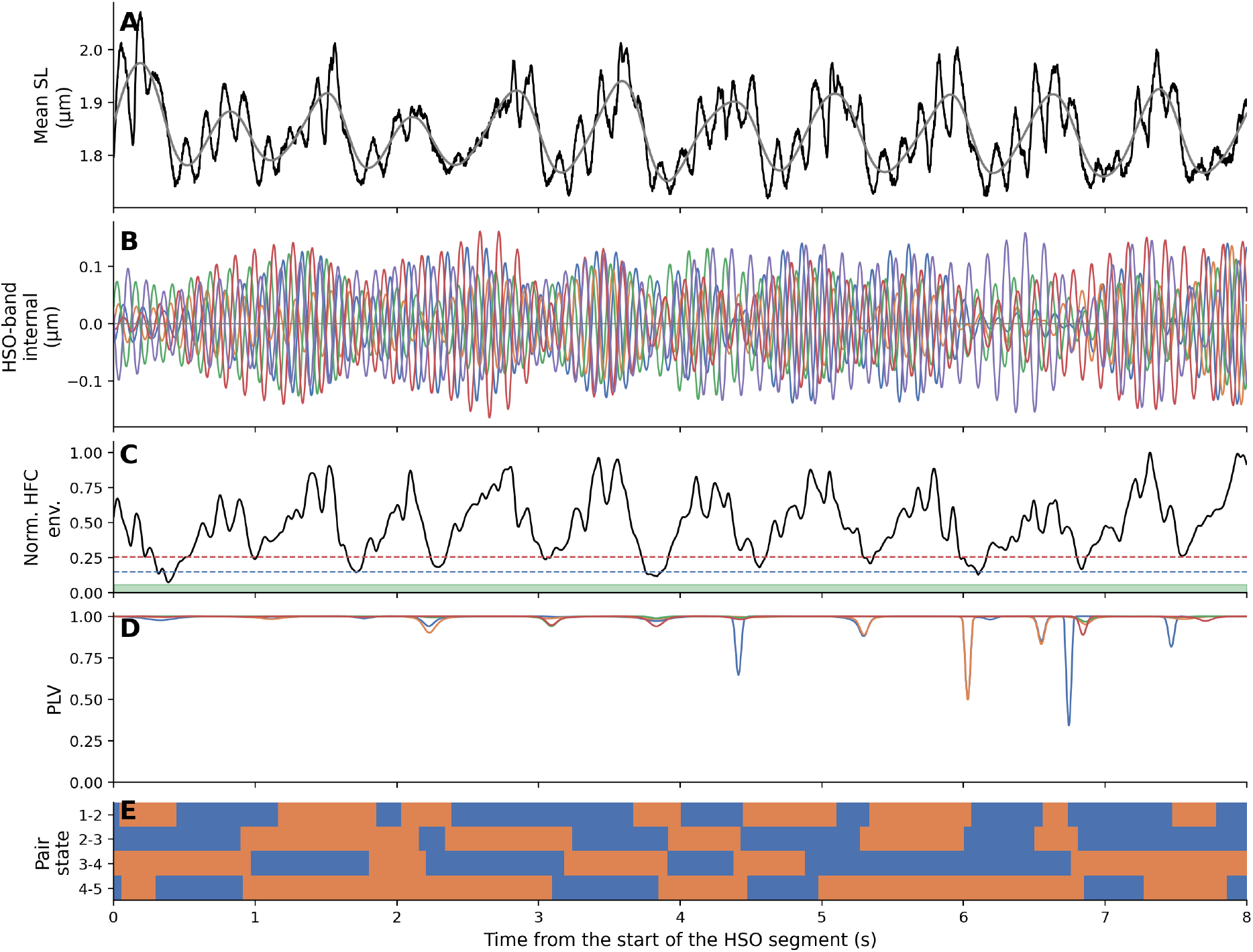
Definition of the local phase state during a representative HSO segment. **A**, local 5-sarcomere mean trace and its low-pass component. **B**, HSO-centered internal signals for the five sarcomeres. **C**, normalized high-frequency envelope and hysteresis thresholds that define the phase-valid gate. **D**, phase-locking value (PLV) for the four neighboring pairs. **E**, binary neighboring-pair states (blue, in-phase; orange, anti-phase) for pairs 1–2, 2–3, 3–4, and 4–5. The segment is shown with the refined visualization gate for display continuity; all quantitative statistics in Figures 3 and 4 use the fixed primary gate.

**Figure 3:**
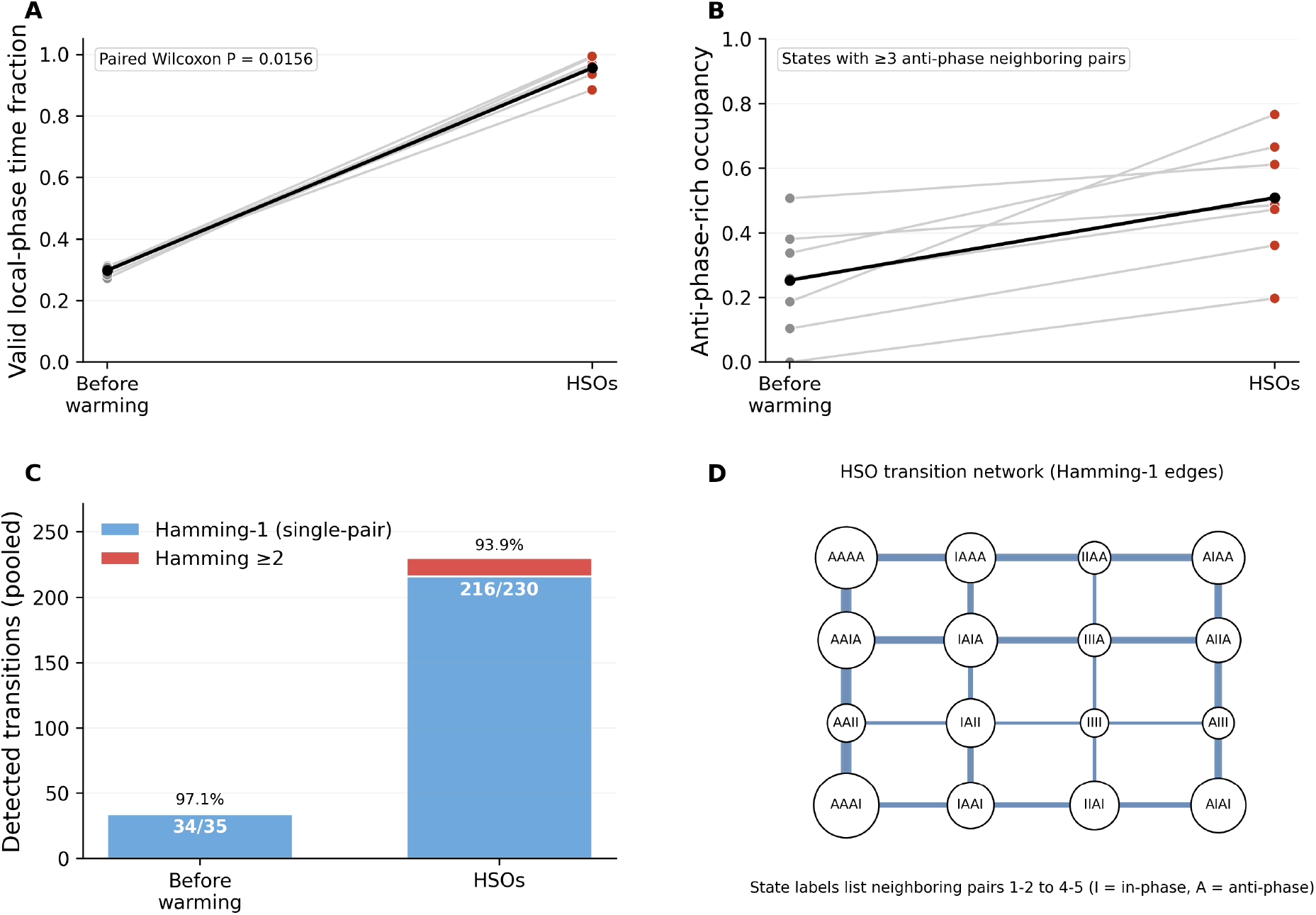
HSOs reveal a constrained local phase topology. **A**, paired values for the fraction of time in which local phase tracking was valid before warming and during HSOs (fixed primary gate). **B**, paired values for time spent in anti-phase-rich states, defined as states with three or more anti-phase neighboring pairs. **C**, pooled transition topology before warming and during HSOs. Numbers inside the bars indicate the count of Hamming-1 transitions relative to all detected transitions. A Hamming-1 transition means that only one of the four neighboring-pair relations changed between successive patterns; along the five-sarcomere chain, this is the smallest possible local update and can be pictured as a one-step move of a local phase boundary. **D**, HSO transition network restricted to Hamming-1 edges. State labels encode the four neighboring pairs from 1–2 to 4–5, with I for in-phase and A for anti-phase. Edge width scales with pooled HSO transition count.

Figure 2 shows how the local phase information was defined in a representative HSO segment. For each cell, the analysis band was centered on the HSO fundamental frequency. The centered-band internal signals, their envelope-derived gate, pairwise phase-locking values, and the four neighboring-pair state traces together show that local phase patterns can be tracked across nearly the entire segment. This continuity makes it possible to analyze trajectories through local state space, not merely isolated pairwise snapshots.

### 2.3 HSOs occupy a constrained local phase topology dominated by single-pair switches

At each valid time point, the five-sarcomere segment was represented by four neighboring-pair relations, each classified as either in-phase or anti-phase. This yields a 16-state local phase network. A Hamming-1 transition means that one neighboring-pair relation changed while the other three remained the same. Along a five-sarcomere line, that is the smallest possible topological update and can be pictured as a one-step movement of a local phase boundary.

Across this 16-state network, transitions were strongly constrained. Under the fixed primary gate, 34 of 35 transitions before warming (97.1%) and 216 of 230 transitions during HSOs (93.9%) were Hamming-1 events (Figure 3C). In every cell, the fraction of Hamming-1 transitions remained high (before warming, 0.875–1.000; HSOs, 0.892–1.000; Supplementary Figure S3A). Thus, local reconfiguration was dominated by one-pair flips rather than by simultaneous multi-pair rearrangements.

This result was stable across parameter variation and gate choice. Across 54 sensitivity settings spanning gate threshold, gap fill, dwell time, and phase-smoothing window, the HSO fraction of Hamming-1 transitions remained between 0.876 and 0.980 (Supplementary Figure S1 and Supplementary Table S2). The HSO transition network (Figure 3D) likewise shows that the most frequent links connect states differing by Hamming distance 1. HSOs therefore define a constrained local phase topology rather than an unstructured local disorder.

### 2.4 HSOs favor anti-phase-rich local configurations

The occupancy of local phase configurations also changed in a structured way. The fraction of valid time spent in states with three or more anti-phase neighboring pairs increased from 0.254 before warming to 0.509 during HSOs (paired Wilcoxon *P* = 0.0156; Figure 3B). The mean number of anti-phase neighboring pairs likewise increased from 1.894 to 2.461 (*P* = 0.0156). HSOs were therefore not only more trackable than the before-warming segment, but were also enriched in local configurations containing multiple anti-phase neighboring-pair relations.

We use the term *anti-phase-rich* for states with three or more anti-phase neighboring pairs because that threshold provided the most balanced descriptor among the alternative occupancy definitions examined in sensitivity analyses (Supplementary Figure S1 and Supplementary Table S3). In other words, the constrained switching topology is accompanied by a shift in state occupancy toward locally anti-phase-rich configurations.

### 2.5 A descriptive cycle-level amplitude-synchrony summary bridges local HSO dynamics and the mean trace

As a complementary analysis of how local coordination appears in the segment-average signal, we constructed cycle-level variables from the trimmed HSO window using the quality-control-valid sarcomere set. Because cell 2 sarcomere 2 was the only trace clearly separated from the others in trimmed-window mean sarcomere length (Supplementary Figure S5A), the cycle-level target was defined on the valid-sarcomere set so that the state variables and averaged signal were defined on the same sarcomeres. For each cycle, *A* was the mean peak-to-peak HSO amplitude across valid sarcomeres, *R_w_* was their weighted synchrony, and *Y*_valid_ was the peak-to-peak HSO amplitude of the valid-sarcomere mean trace (Figure 4A).

**Figure 4:**
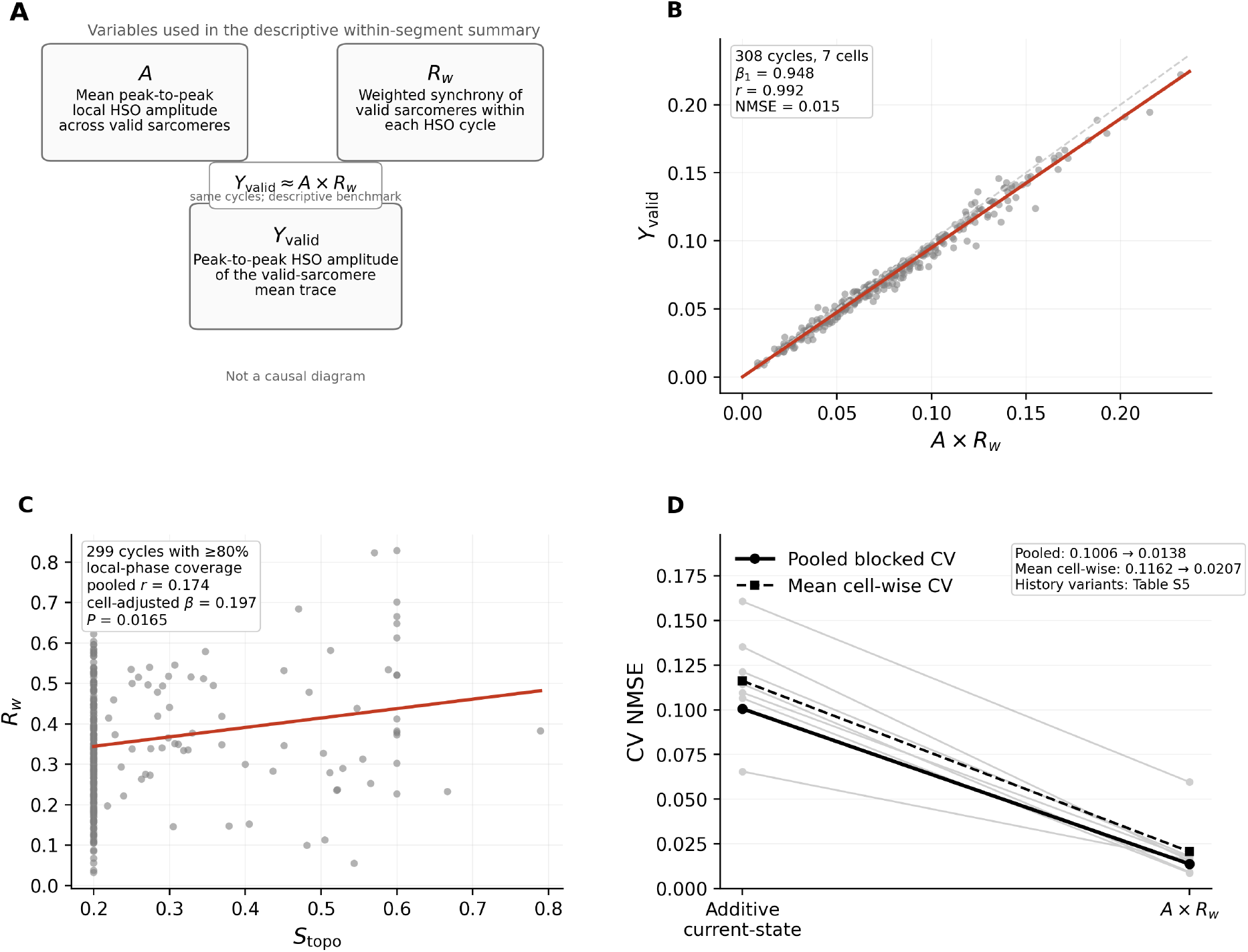
A descriptive amplitude-synchrony summary bridges local HSO dynamics and the mean trace. **A**, schematic definition of the cycle-level variables used in the descriptive within-segment summary. This panel is not intended as a causal diagram. *Y*_valid_ is the peak-to-peak HSO amplitude of the mean trace formed from the quality-control-valid sarcomeres, *A* is the mean peak-to-peak HSO amplitude across those sarcomeres, and *R_w_* is their weighted synchrony. The trimmed window and valid-sarcomere set are described in Methods. **B**, pooled relation between *Y*_valid_ and *A × R_w_* across 308 cycles from seven cells; dashed line, identity; red line, pooled direct fit. Because all three quantities are derived from the same observed fast-component signals in the same cycles, this panel is interpreted as a descriptive within-segment summary rather than as a standalone cross-segment prediction rule. **C**, bridge from the Figure 3 local-state topology to the continuous synchrony term used in the cycle-level analysis. *S*_topo_ is a cycle-average state-derived synchrony factor computed from the 16 local phase patterns under an equal-amplitude 0*/π* assignment; only cycles with at least 80% local-phase coverage were included. The red line is the pooled linear fit, and the annotation reports the modest cell-adjusted association with *R_w_*. **D**, blocked cross-validation comparison between an additive current-state model and the *A × R_w_* summary; gray lines, individual cells; black solid line, pooled performance; black dashed line, mean cell-wise performance. Detailed cell-wise slopes and history-augmented models are summarized in Supplementary Table S5.

To create a minimal bridge between the binary local-state description of Figure 3 and the continuous synchrony term used below, we assigned each of the 16 local phase patterns a state-derived synchrony factor *S*_topo_ = *|* ∑ *_k_ s_k_|/*5 under the equal-amplitude, 0*/π* phase assignment implied by its four neighboring-pair relations, and averaged *S*_topo_ within each HSO cycle. *S*_topo_ was lower in cycles that spent more time in anti-phase-rich states (pooled *r* = −0.402) and showed a modest positive association with *R_w_* among cycles with at least 80% local-phase coverage (299 cycles; pooled *r* = 0.174; cell-adjusted *β* = 0.197, *P* = 0.0165; Figure 4C). These results do not imply a one-to-one mapping from binary local states to *R_w_*, but they do show that cycles with more synchronized local phase patterns tend to have higher *R_w_*. The topology quantified in Figure 3 therefore connects naturally to the synchrony summary used in the cycle-level analysis.

Across 308 cycles from seven cells, *Y*_valid_ was closely approximated by the product *A × R_w_* (Figure 4B). A direct pooled fit yielded *β*_1_ = 0.948, *β*_0_ *≈* 0, *r* = 0.992, and normalized mean squared error 0.015. Cell-wise slopes were likewise close to 1 (0.908–0.972; Supplementary Table S5), indicating that this approximation was not created by one outlier cell. Because *Y*_valid_, *A*, and *R_w_* are all computed from the same observed fast-component signals within the same cycle, we interpret this result as a descriptive decomposition of the observed mean signal rather than as a standalone cross-segment prediction rule.

We then asked whether the same observed mean signal could be summarized equally well by a more generic additive current-state model or whether simple history terms materially improved the *A × R_w_* summary. In blocked cross-validation, the *A × R_w_* summary strongly outperformed the additive model (pooled normalized mean squared error 0.0138 vs 0.1006; mean cell-wise normalized mean squared error 0.0207 vs 0.1162; Figure 4D). Adding *Y*_valid,*n*−1_, (*A × R_w_*)_*n*−1_, or a two-step history summary changed pooled error only marginally (0.0135–0.0137; Supplementary Table S5). Accordingly, within the same observed segment, a simple product of local amplitude and synchrony provides a parsimonious cycle-level summary of the mean fast HSO amplitude, with little additional contribution from these simple history terms.

## 3 Discussion

The present reanalysis clarifies two connected aspects of HSO coordination in living cardiomyocytes. The main result is that neighboring sarcomeres reconfigured through a constrained 16-state phase topology dominated by Hamming-1 transitions and accompanied by increased anti-phase-rich occupancy. In the same segments, the HSO amplitude of the valid-sarcomere mean trace was closely approximated by the product of local amplitude and weighted synchrony as a complementary within-segment summary. Together, these analyses argue against unstructured local disorder and instead define HSOs as a structured neighboring-sarcomere rephasing regime.

This finding adds a temporal and topological measure to the broader literature on sarcomere heterogeneity. Stretch-dependent harmonization of sarcomere strain (Li et al., 2023), myofibrillar variation in sarcomere length (Lookin et al., 2023), and in vivo evidence that sarcomeric synchrony affects ventricular pump performance (Kobirumaki-Shimozawa et al., 2021) all indicate that local nonuniformity is part of cardiac function rather than an experimental nuisance. Recent work on living myocardium further emphasizes that asynchronous sarcomere movement persists under physiological conditions and may reflect mechanically meaningful differences between neighboring elements (Kobirumaki-Shimozawa et al., 2024). The present analysis extends that view by showing how local phase relations among adjacent sarcomeres are reconfigured over time and how the resulting local coordination is reflected in the mean HSO signal.

The predominance of Hamming-1 transitions fits naturally with prior work on inter-sarcomere coordination in oscillatory or perturbed contractile systems. Quick-stretch experiments revealed adjacent clusters of yielding sarcomeres (Shimamoto et al., 2009), phosphate-jump experiments identified sarcomere “give” that starts locally and propagates to neighbors (Stehle, 2017), and SPOC studies have long emphasized succession of lengthening phases and wave-like coordination across neighboring contractile elements (Yasuda et al., 1996; Sasaki et al., 2005; Kagemoto et al., 2015). The present recordings span only five consecutive sarcomeres along one line, so they do not reconstruct a propagating wave directly. Even so, the observed topology is consistent with a nearest-neighbor-like mode of reconfiguration in which local phase boundaries shift stepwise rather than the pattern being reset wholesale. Such a regime could allow local phase lags or staggered relaxation to develop while beat-scale timing remains comparatively stable.

The cycle-level amplitude-synchrony summary adds a complementary quantitative interpretation. The mean HSO amplitude of the valid-sarcomere average was closely approximated by *A × R_w_*, where *A* describes how strongly the local sarcomeres oscillate and *R_w_* describes how efficiently those rapid motions survive averaging. Because *Y*_valid_, *A*, and *R_w_* are all constructed from the same observed fast-component signals within the same cycles, this should not be read as a first-principles derivation or as a standalone law-like prediction rule. Rather, it is a benchmark decomposition of the observed mean signal into amplitude and synchrony terms within the same segment, consistent with weighted phasor-like averaging of narrowband components. In that descriptive sense, the gap between the additive model and the *A × R_w_* summary is informative: it indicates that a multiplicative amplitude-synchrony summary is markedly more parsimonious for this dataset, whereas simple history terms contribute little additional explanatory value.

The topology and cycle-level analyses still live at different descriptive levels, and the present data do not justify a one-to-one mapping from a binary local state to the continuous synchrony variable *R_w_*. Even so, the new state-derived synchrony factor provides a useful bridge between them. Because *S*_topo_ is computed directly from the 16 local phase patterns, its positive association with *R_w_* indicates that the Figure 3 topology carries the same synchrony directionality used in Figure 4, without implying that anti-phase-rich occupancy alone determines the measured mean amplitude. The local topology analysis therefore identifies the constrained neighboring-sarcomere reconfiguration, whereas the cycle-level analysis shows how local amplitude and synchrony provide a compact descriptive summary of the mean signal seen in the same segment. An earlier exploratory attenuation/cancellation analysis from the same dataset is retained in the Supplementary Information as supportive material rather than as a main-text pillar.

This hierarchy of claims is intentional. The strongest findings are the gain in phase-trackable time during HSOs, the dominance of Hamming-1 local updates, and the shift toward anti-phase-rich occupancy. The *A × R_w_* analysis is presented one step below these results, as a descriptive bridge from local coordination to the observed mean fast signal in the same segment. Framing the results this way reduces over-interpretation while preserving the biological message that local HSO nonuniformity is structured rather than random.

The warmed-cell context helps place these results in a physiologically interpretable framework. Thermal activation of thin filaments can bias the contractile system toward activation even without large Ca^2+^ changes (Oyama et al., 2012; Ishii et al., 2019, 2020). At the same time, cardiac activation is shaped by cooperative control involving cross-bridges, titin-associated effects, and thick-filament regulatory states (Solís and Solaro, 2021; Park-Holohan et al., 2021). Length- and strain-dependent activation likewise show that local mechanics and activation are tightly coupled (de Tombe et al., 2010; Campbell, 2011). Within that framework, minimal single-pair switching offers a plausible way for neighboring sarcomeres to accommodate local phase offsets without losing beat-scale timing, whereas the amplitude-synchrony summary provides an explicit within-segment description of how those local dynamics become visible in the segment-average oscillatory signal. The molecular mechanism of the switches and synchrony changes remains unresolved, but the organization of the phenomenon is now better defined.

From a biophysical perspective, this matters because it identifies an experimentally tractable mesoscale coordination unit between single-sarcomere heterogeneity and the averaged signals by which beating cells are usually judged. From a physiological perspective, it suggests that local nonuniformity during HSOs need not be read as mere disorder, but can instead reflect structured redistribution of timing within a still-beating cardiomyocyte. A recent follow-up reanalysis of the same recordings suggests that this constrained regime can also support a continuous ordering of local coordination states and rankable mesoscale readouts, reinforcing the view that the present topology is a foundation rather than an endpoint description of HSO coordination (Shintani, 2026a).

This study also has clear limitations. It analyzes seven cells and one five-sarcomere line segment from each cell. The primary topology analysis requires the intact five-sarcomere chain, whereas the cycle-level analysis uses the quality-control-valid sarcomere set so that the state variables and averaged target are defined on the same sarcomeres. The present work therefore offers a quantitative working description rather than a final general law. Independent recordings, additional temperature conditions, longer line segments, and simultaneous measurements of local Ca^2+^, force, or titin-related mechanics will be needed to test generality and mechanism directly. Simultaneous imaging of local Ca^2+^ and single-sarcomere dynamics has already been achieved in neonatal cardiomyocytes (Tsukamoto et al., 2016), and recent in vivo work connects sarcomeric synchrony and asynchrony to ventricular mechanics (Kobirumaki-Shimozawa et al., 2021, 2024). The present dataset therefore provides a tractable spatial and quantitative unit for those next experiments.

During HSOs, neighboring sarcomeres do not wander through local phase space at random. They reconfigure through a constrained local phase topology dominated by minimal single-pair switches, and the mean oscillatory amplitude observed in the same segment is parsimoniously summarized by a cycle-level amplitude-synchrony summary. This view places the HSO state on firmer quantitative footing by linking local sarcomere coordination to averaged measurements in living cardiomyocytes while keeping the hierarchy of claims explicit.

## 4 Materials and methods

### 4.1 Study design, data source, and ethics

This study is a reanalysis of previously acquired sarcomere-length data from neonatal rat cardiomyocytes. No new animal experiments were performed for the present work. The original experimental procedures were approved by the Animal Experiment Committee of the Faculty of Science at the University of Tokyo and were conducted according to the Animal Experiment Implementation Manual of the University of Tokyo, as reported previously (Shintani et al., 2020). The reanalyzed dataset consisted of one time series and data from seven cells, with five consecutive sarcomere-length traces analyzed for each cell.

### 4.2 Original cell preparation and microscopic system

The original experimental procedures have been described in detail elsewhere (Shintani et al., 2014, 2015, 2020). Briefly, immature cardiomyocytes were isolated from 1-day-old Wistar rats and cultured in DMEM/F-12 containing 10% fetal bovine serum, 100 U/ml penicillin, and 100 U/ml streptomycin. Cells were transfected with pAcGFP-actinin, and measurements were performed 1 day after transfection. Imaging used an inverted microscope equipped with a high-sensitivity camera and an oil-immersion objective. The pixel size in the specimen plane was 150 nm, and the frame rate was 500 frames/s for high-temporal-resolution analysis. A 1550-nm laser was used as a local heat source, and a 488-nm laser was used to excite AcGFP-actinin fluorescence (Oyama et al., 2012; Shintani et al., 2020).

### 4.3 Sarcomere-length measurement and segment definition

Sarcomere length was measured by fitting the *α*-actinin-AcGFP intensity profile at each Z line to a parabolic function, as described previously (Shintani et al., 2014). The standard deviation of the sarcomere-length estimate was approximately 4 nm at 500 Hz (Shintani et al., 2020). The present reanalysis used seven cells, each represented by five consecutive sarcomeres. For the primary neighboring-pair topology analysis, the intact five-sarcomere chain was retained. Non-topological summaries were additionally checked in sensitivity analyses. For the complementary cycle-level analysis, one sarcomere trace (cell 2, sarcomere 2) was excluded from the quality-controlled sarcomere set because it was the only trace clearly separated from the others in trimmed-window mean sarcomere length; the state variables and the averaged target were then defined on the same sarcomeres.

### 4.4 Signal decomposition and cell-specific HSO-centered phase tracking

For each cell, the HSO fundamental frequency was estimated from the HSO segment by Welch spectral analysis of the root-mean-square internal signal over 4.5–9.5 Hz. A cell-specific centered analysis band was then defined as *f*_0,HSO_ *±* 1.0 Hz, clipped to the 3.5–25 Hz high-frequency band. To isolate internal sarcomeric motion, each sarcomere trace was centered by subtracting the instantaneous local mean of the five sarcomeres and then band-pass filtered in the HSO-centered band. Instantaneous phase and analytic amplitude envelope were obtained from the Hilbert transform.

Phase-valid intervals were defined from the mean high-frequency envelope using a hysteresis gate. The fixed primary gate used an upper quantile threshold of 0.46, a lower-threshold offset of 0.12, a minimum on-duration of 0.05 s, and a gap-fill duration of 0.03 s. A refined visualization gate with the same thresholds but a gap-fill duration of 0.12 s was used only for display continuity in Figure 2 and supplementary sensitivity analyses.

### 4.5 Neighboring-pair states and transition topology

For each of the four neighboring sarcomere pairs (1–2, 2–3, 3–4, and 4–5), phase differences were smoothed in a sliding window corresponding to approximately one-half HSO cycle. A neighboring pair was classified as in-phase when cos(Δ*ϕ*) ≥ 0 and anti-phase when cos(Δ*ϕ*) *<* 0. The four binary neighboring-pair states yielded 16 possible local phase patterns. State transitions were detected after enforcing a minimum dwell time of 0.02 s and a maximum gap of 0.04 s. The Hamming distance between consecutive states was used to classify whether a transition involved a change in one neighboring pair or in two or more neighboring pairs. To create a minimal bridge from the binary local-state description to the continuous synchrony term used in the cycle-level analysis, each 16-state pattern was also assigned a state-derived synchrony factor

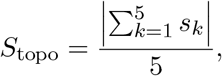

where *s_k_* ∈ {+1, −1} denotes the equal-amplitude, 0*/π* phase assignment implied by the neighboring-pair pattern. This equal-amplitude construction was used only as a coarse bridge from the binary local-state description to the measured synchrony term and was not interpreted as a one-to-one map to the continuous synchrony observed in the data. Cycle-mean *S*_topo_ was computed within HSO cycles using only cycles with at least 80% local-phase coverage.

### 4.6 Occupancy metrics and exploratory attenuation/cancellation analyses

The principal occupancy metric used in the main text was the fraction of valid time spent in states with three or more anti-phase neighboring pairs. We refer to these as *anti-phase-rich* states. Beat minima and maxima were identified from the low-pass local 5-sarcomere mean trace using prominence-based peak detection with a minimum inter-minimum distance of 0.5 s. Beat-level intervals between successive minima were used only for exploratory supplementary analyses.

For those supplementary analyses, the local 5-sarcomere mean-trace high-frequency attenuation index (attenuation index) was defined as

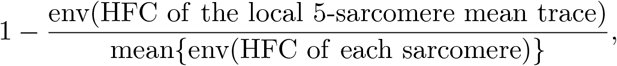

where env(·) denotes the Hilbert amplitude envelope. Higher values indicate greater attenuation of the high-frequency component in the local mean relative to the individual sarcomere traces. We also defined an exploratory phase-vector cancellation index from the HSO-centered analytic signals *z_k_*(*t*) of the five sarcomeres:

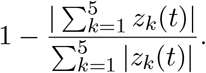

This quantity was averaged over valid time points within each beat and is reported only in the Supplementary Information.

### 4.7 Complementary cycle-level amplitude-synchrony analysis

For the complementary cycle-level analysis, the HSO window was trimmed by 0.25 s at each boundary (9.75–17.25 s) and downsampled fivefold to approximately 100 Hz. Valid sarcomeres were defined as all traces except cell 2 sarcomere 2, which was treated as a quality-control outlier because it was the only trace clearly separated from the others in trimmed-window mean sarcomere length. Valid and all-sarcomere traces were band-pass filtered at 4–12 Hz to isolate the fast HSO component. The low-frequency phase covariate was derived from a 0.5–2.0 Hz band-pass filter applied to the valid-sarcomere mean trace.

Cycles were identified from the unwrapped Hilbert phase of the fast component of the valid-sarcomere mean trace and retained when their duration was 0.08–0.25 s. For each cycle, local amplitude *A* was the mean peak-to-peak fast-component amplitude across valid sarcomeres. The canonical target *Y*_valid_ was the peak-to-peak fast-component amplitude of the valid-sarcomere mean trace. Weighted synchrony *R_w_* was the cycle-average of

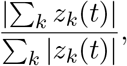

computed from the analytic fast-component signals *z_k_*(*t*) of the valid sarcomeres. Because *Y*_valid_, *A*, and *R_w_* are all computed from the same analytic fast-component signals within the same cycles, the product form *A × R_w_* was interpreted here as a descriptive approximation rather than as an independent law. In narrowband analytic signals, the magnitude of the cycle-wise segment mean naturally depends on both an overall amplitude scale and the degree of phase alignment among components. Under that phasor-like averaging picture, a product of an amplitude summary and a synchrony summary is therefore expected to provide a compact within-segment approximation to the observed mean fast amplitude. This interpretation does not constitute a first-principles derivation and is not assumed to generalize beyond the present observed segment without further validation. A second-order synchrony summary *R*_2_ was computed analogously from 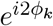 and was used only in additive comparison models. The all-sarcomere mean target *Y*_all_ was retained only for descriptive comparison and supplementary target-sensitivity analyses. It was not used as the canonical target because it includes the quality-control-outlier trace and is therefore not defined on the same sarcomere set as the valid-sarcomere state variables.

### 4.8 Statistics and additional analyses

Paired before-versus-HSO comparisons across cells were summarized by means, paired mean differences, 95% confidence intervals, Wilcoxon signed-rank *P* values, and sign-test *P* values. Direct cycle-level fits used ordinary least squares for *Y*_valid_ ∼ *β*_0_ + *β*_1_(*A × R_w_*) as a descriptive within-segment benchmark fit. Model comparison for the cycle-level amplitude-synchrony analysis used ridge regression (*α* = 10^−3^) with blocked fivefold cross-validation within each cell and in pooled cell-blocked folds. Candidate models were an additive current-state model (*A, R_w_, R*_2_, and slow-phase nuisance terms), *A × R_w_, A × R_w_* + *Y*_valid,*n*−1_, *A × R_w_* + (*A × R_w_*)_*n*−1_, and *A × R_w_* plus a two-step history summary. Equivalent direct-fit and blocked-cross-validation comparisons were repeated for *Y*_all_ only as supplementary target-sensitivity analyses. Because *Y*_valid_, *A*, and *R_w_* are derived from the same observed fast-component signals, these direct-fit and blocked-cross-validation analyses were interpreted as tests of descriptive parsimony within the same segment rather than of independent predictive generality. Associations between cycle-mean *S*_topo_ and *R_w_* were summarized by pooled correlation and by ordinary least squares with cell fixed effects and HC3 standard errors. Permutation tests with 500 permutations assessed improvement in pooled blocked-cross-validation error when comparing nested models. Because history terms produced only very small absolute changes in error, practical magnitude rather than *P* value alone was emphasized in interpretation.

Exploratory beat-phase, attenuation/cancellation, and frequency-dispersion analyses are summarized in the Supplementary Information. Parameter robustness of the Hamming-1 transition dominance and of the exploratory attenuation association was assessed across 54 settings spanning gate threshold, gap-fill duration, dwell time, and phase-smoothing window.

## Author contributions

S.A.S. conceived the study, performed the original experiments and the present reanalysis, and wrote the manuscript.

## Data and code availability

The reanalyzed dataset, the figure-level summary tables used for the present analyses, and the custom analysis code used in this study are available from the corresponding author upon reasonable request. Selected source-data tables and figure-generation materials used during manuscript preparation are organized in the accompanying revision package for transparency and reproducibility.

## Declaration of generative AI and AI-assisted technologies in the writing process

During the preparation of this work, the author used ChatGPT (OpenAI) to improve the readability and language of the manuscript. After using this tool, the author reviewed and edited the content as needed and takes full responsibility for the content of the publication.

## References

Garg, A., K.J. Lavine, and M.J. Greenberg. 2024. Assessing cardiac contractility from single molecules to whole hearts. JACC Basic Transl. Sci. 9:414–439. doi: 10.1016/j.jacbts.2023.07.013.

Rassier, D.E., and A. Månsson. 2025. Mechanisms of myosin II force generation: insights from novel experimental techniques and approaches. Physiol. Rev. 105:1–93. doi: 10.1152/physrev.00014.2023.

Li, J., J. Sundnes, Y. Hou, M. Laasmaa, M. Ruud, A. Unger, T.R. Kolstad, M. Frisk, P.A. Norseng, L. Yang, I.E. Setterberg, E.S. Alves, M. Kalakoutis, O.M. Sejersted, J.T. Lanner, W.A. Linke, I.G. Lunde, P.P. de Tombe, and W.E. Louch. 2023. Stretch harmonizes sarcomere strain across the cardiomyocyte. Circ. Res. 133:255–270. doi: 10.1161/CIRCRESAHA.123.322588.

Lookin, O., P. de Tombe, N. Boulali, C. Gergely, T. Cloitre, and O. Cazorla. 2023. Cardiomyocyte sarcomere length variability: membrane fluorescence versus second harmonic generation myosin imaging. J. Gen. Physiol. 155:e202213289. doi: 10.1085/jgp.202213289.

Kobirumaki-Shimozawa, F., T. Shimozawa, K. Oyama, S. Baba, J. Li, T. Nakanishi, T. Terui, W.E. Louch, Shin’ichi Ishiwata, and N. Fukuda. 2021. Synchrony of sarcomeric movement regulates left ventricular pump function in the in vivo beating mouse heart. J. Gen. Physiol. 153:e202012860. doi: 10.1085/jgp.202012860.

Kobirumaki-Shimozawa, F., K. Oyama, T. Nakanishi, Shin’ichi Ishiwata, and N. Fukuda. 2024. Asynchronous movement of sarcomeres in myocardium under living conditions: role of titin. Front. Physiol. 15:1426545. doi: 10.3389/fphys.2024.1426545.

Fabiato, A., and F. Fabiato. 1978. Myofilament-generated tension oscillations during partial calcium activation and activation dependence of the sarcomere length–tension relation of skinned cardiac cells. J. Gen. Physiol. 72:667–699. doi: 10.1085/jgp.72.5.667.

Linke, W.A., M.L. Bartoo, and G.H. Pollack. 1993. Spontaneous sarcomeric oscillations at intermediate activation levels in single isolated cardiac myofibrils. Circ. Res. 73:724–734. doi: 10.1161/01.RES.73.4.724.

Yasuda, K., Y. Shindo, and S. Ishiwata. 1996. Synchronous behavior of spontaneous oscillations of sarcomeres in skeletal myofibrils under isotonic conditions. Biophys. J. 70:1823–1829. doi: 10.1016/S0006-3495(96)79747-3.

Shimamoto, Y., M. Suzuki, S.V. Mikhailenko, K. Yasuda, and S. Ishiwata. 2009. Inter-sarcomere coordination in muscle revealed through individual sarcomere response to quick stretch. Proc. Natl. Acad. Sci. USA 106:11954–11959. doi: 10.1073/pnas.0813288106.

Stehle, R. 2017. Force responses and sarcomere dynamics of cardiac myofibrils induced by rapid changes in [Pi]. Biophys. J. 112:356–367. doi: 10.1016/j.bpj.2016.11.005.

Sasaki, D., H. Fujita, N. Fukuda, S. Kurihara, and S. Ishiwata. 2005. Auto-oscillations of skinned myocardium correlating with heartbeat. J. Muscle Res. Cell Motil. 26:93–101. doi: 10.1007/s10974-005-0249-2.

Kagemoto, T., A. Li, C. dos Remedios, and S. Ishiwata. 2015. Spontaneous oscillatory contraction (SPOC) in cardiomyocytes. Biophys. Rev. 7:15–24. doi: 10.1007/s12551-015-0165-7.

Oyama, K., A. Mizuno, S.A. Shintani, H. Itoh, T. Serizawa, N. Fukuda, M. Suzuki, and Shin’ichi Ishiwata. 2012. Microscopic heat pulses induce contraction of cardiomyocytes without calcium transients. Biochem. Biophys. Res. Commun. 417:607–612. doi: 10.1016/j.bbrc.2011.12.015.

Ishii, S., K. Oyama, T. Arai, H. Itoh, S.A. Shintani, M. Suzuki, F. Kobirumaki-Shimozawa, T. Terui, N. Fukuda, and Shin’ichi Ishiwata. 2019. Microscopic heat pulses activate cardiac thin filaments. J. Gen. Physiol. 151:860–869. doi: 10.1085/jgp.201812243.

Ishii, S., K. Oyama, S.A. Shintani, F. Kobirumaki-Shimozawa, Shin’ichi Ishiwata, and N. Fukuda. 2020. Thermal activation of thin filaments in striated muscle. Front. Physiol. 11:278. doi: 10.3389/fphys.2020.00278.

Solís, C., and R.J. Solaro. 2021. Novel insights into sarcomere regulatory systems control of cardiac thin filament activation. J. Gen. Physiol. 153:e202012777. doi: 10.1085/jgp.202012777.

Park-Holohan, S.-J., E. Brunello, T. Kampourakis, M. Rees, M. Irving, and L. Fusi. 2021. Stress-dependent activation of myosin in the heart requires thin filament activation and thick filament mechanosensing. Proc. Natl. Acad. Sci. USA 118:e2023706118. doi: 10.1073/pnas.2023706118.

de Tombe, P.P., R.D. Mateja, K. Tachampa, Y.A. Mou, G.P. Farman, and T.C. Irving. 2010. Myofilament length dependent activation. J. Mol. Cell. Cardiol. 48:851–858. doi: 10.1016/j.yjmcc.2009.12.017.

Campbell, K.S. 2011. Impact of myocyte strain on cardiac myofilament activation. Pflügers Arch. Eur. J. Physiol. 462:3–14. doi: 10.1007/s00424-011-0952-3.

Shintani, S.A., K. Oyama, N. Fukuda, and Shin’ichi Ishiwata. 2015. High-frequency sarcomeric auto-oscillations induced by heating in living neonatal cardiomyocytes of the rat. Biochem. Biophys. Res. Commun. 457:165–170. doi: 10.1016/j.bbrc.2014.12.077.

Shintani, S.A., T. Washio, and H. Higuchi. 2020. Mechanism of contraction rhythm homeostasis for hyperthermal sarcomeric oscillations of neonatal cardiomyocytes. Sci. Rep. 10:20468. doi: 10.1038/s41598-020-77443-x.

Shintani, S.A. 2022. Hyperthermal sarcomeric oscillations generated in warmed cardiomyocytes control amplitudes with chaotic properties while keeping cycles constant. Biochem. Biophys. Res. Commun. 611:8–13. doi: 10.1016/j.bbrc.2022.04.055.

Shintani, S.A. 2025. Chaordic homeodynamics: the periodic chaos phenomenon observed at the sarcomere level and its physiological significance. Biochem. Biophys. Res. Commun. 760:151712. doi: 10.1016/j.bbrc.2025.151712.

Tsukamoto, S., T. Fujii, K. Oyama, S.A. Shintani, T. Shimozawa, F. Kobirumaki-Shimozawa, Shin’ichi Ishiwata, and N. Fukuda. 2016. Simultaneous imaging of local calcium and single sarcomere length in rat neonatal cardiomyocytes using yellow Cameleon-Nano140. J. Gen. Physiol. 148:341–355. doi: 10.1085/jgp.201611604.

Shintani, S.A., K. Oyama, F. Kobirumaki-Shimozawa, T. Ohki, Shin’ichi Ishiwata, and N. Fukuda. 2014. Sarcomere length nanometry in rat neonatal cardiomyocytes expressed with α-actinin-AcGFP in Z discs. J. Gen. Physiol. 143:513–524. doi: 10.1085/jgp.201311118.

Shintani, S.A. 2026a. A shared rephasing compass reveals structured local mismatch placement during hyperthermal sarcomeric oscillations. bioRxiv. doi: 10.64898/2026.03.26.714639.

