## Supplementary Information for "Constrained neighboring-sarcomere phase topology relates to mean HSO amplitude in living cardiomyocytes"

Seine A. Shintani

#### Supplementary Note

This supplement is organized around the same claim hierarchy used in the revised main manuscript and is intended to document robustness, scope, and limits without elevating exploratory analyses above the topology-centered main results. The *primary* claim is the constrained local phase topology during HSOs, supported by three main-text pillars: greatly increased valid time for local phase tracking, dominance of Hamming-1 transitions, and increased occupancy of anti-phase-rich states. The *descriptive bridge* claim is the cycle-level $A \times R_w$ summary for the mean fast signal within the same observed segment. *Supportive or exploratory* material includes beat-phase displays and the earlier attenuation/cancellation analyses, which are retained for transparency but are not treated as co-equal main results.

Accordingly, Supplementary Figure S1 separates robustness of the primary topology pillars from robustness of the supportive attenuation association. Supplementary Figure S2 provides within-cell null controls for that supportive attenuation analysis. Supplementary Figure S3 combines a cell-wise display of the primary Hamming-1 result with the beat-phase partition used only for exploratory analyses. Supplementary Figure S4 retains the earlier attenuation/cancellation figure explicitly as supportive material after the manuscript was refocused around constrained local phase topology as the main claim and the cycle-level amplitude-synchrony summary as a complementary within-segment bridge. Supplementary Figure S5 documents the target-choice and parsimony checks for the descriptive bridge by visualizing the only trace clearly separated from the others in trimmed-window mean sarcomere length, comparing $Y_{\text{valid}}$ with $Y_{\text{all}}$ in direct fits and blocked cross-validation, and showing that the apparent gain from a simple history term is inflated when that trace is allowed back into the target. Supplementary Table S5 expands the descriptive-bridge statistics to include the additive model, the $A \times R_w$ summary, and the three history-augmented variants on both pooled and cell-wise bases. Exploratory analyses that did not receive sufficient support from the present dataset are listed explicitly in Supplementary Table S4 so that the scope and limits of the present evidence remain clear.

#### Supplementary Figure S1

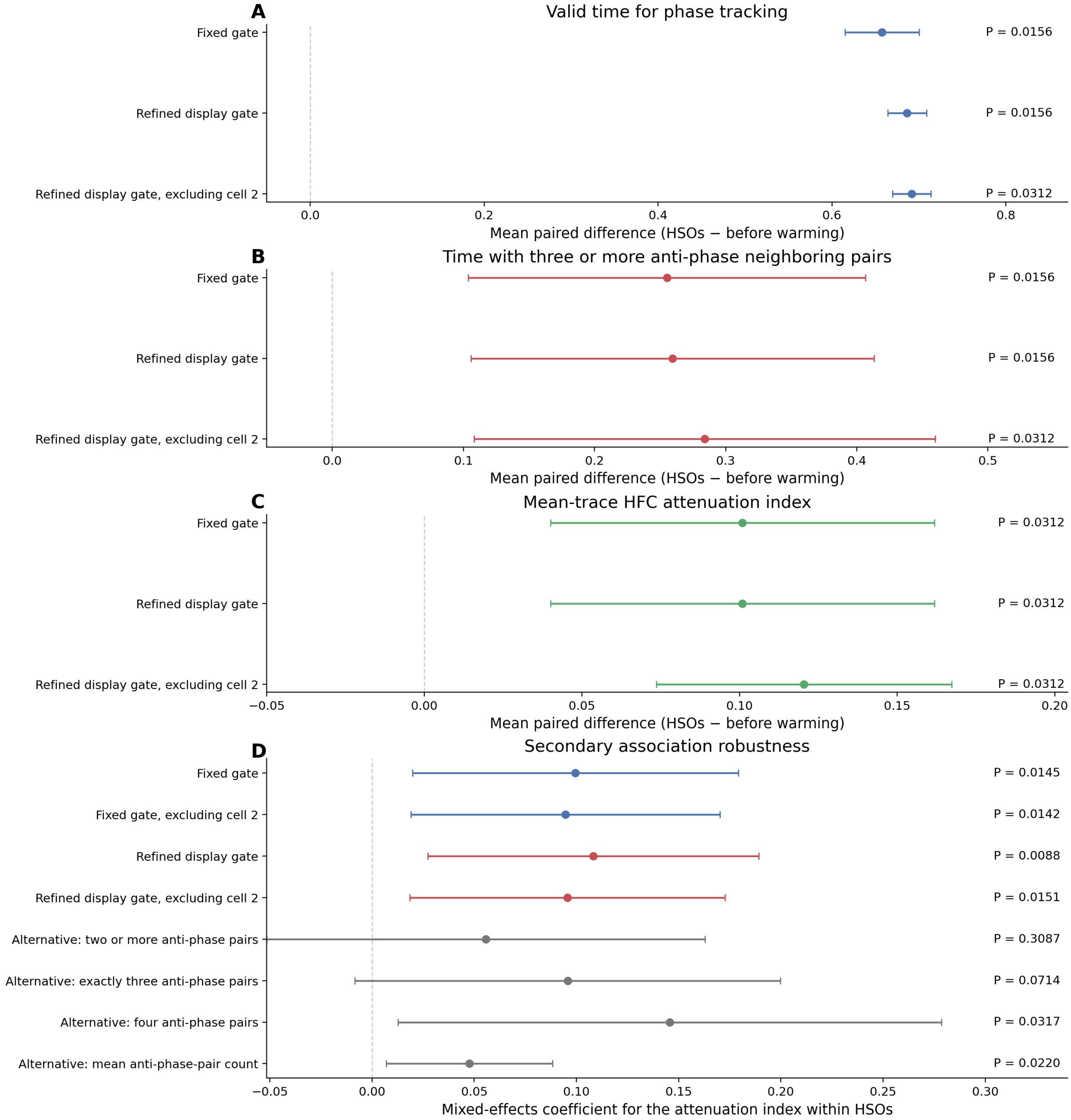


**Robustness of the primary topology pillars and of the supportive HSO beat-level attenuation association. A–B**, mean paired differences between HSOs and before warming for valid time for phase tracking and time with three or more anti-phase neighboring pairs. These are direct robustness checks for two of the three main-text primary pillars. **C**, mean paired difference for the local 5-sarcomere mean-trace high-frequency attenuation index, retained as supportive rather than primary. Results are shown for the fixed primary gate, the refined display gate, and the refined display gate after exclusion of cell 2. Error bars indicate the 95% confidence interval of the paired difference across cells. **D**, robustness of the beat-level attenuation association under gate choice, exclusion of cell 2, and alternative anti-phase occupancy definitions. The positive association remained for the fixed and refined gates and after exclusion of cell 2; alternative occupancy definitions were more variable, reinforcing the decision to treat this analysis as supportive rather than as a main-text pillar.

### Supplementary Figure S2

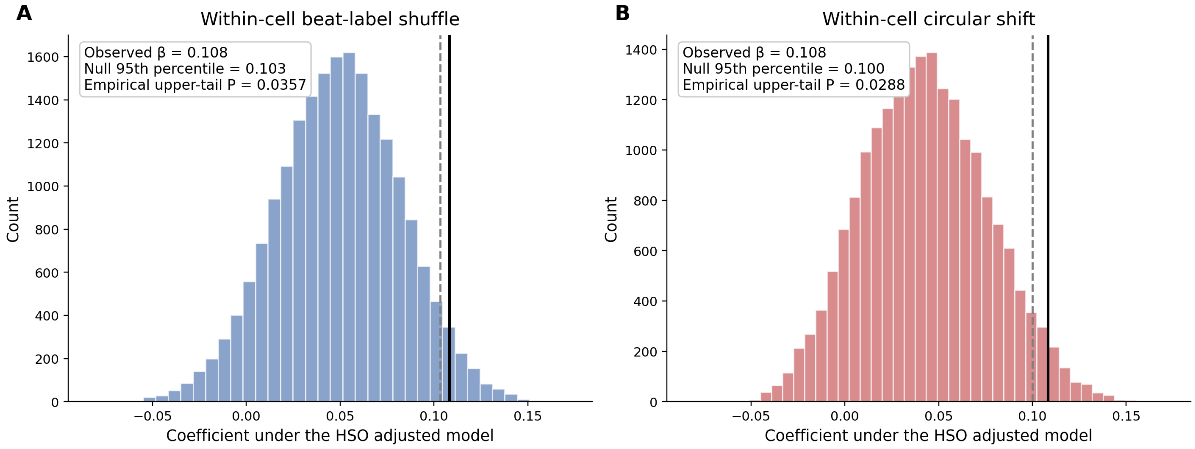


**Within-cell null controls for the supportive HSO beat-level attenuation association.** Null distributions of the adjusted HSO coefficient were generated by perturbing the beat-wise anti-phase-rich predictor within each cell while preserving the observed beat duration and transition-density covariates. In the beat-label shuffle null, predictor values were randomly permuted within each cell. In the circular-shift null, the predictor sequence was shifted within each cell by a non-zero random offset. The black solid line marks the observed coefficient and the gray dashed line marks the 95th percentile of the null distribution. These null controls are provided to document that the supportive association is not obviously produced by simple within-cell rearrangements, while still remaining secondary to the topology-centered main claim.

### Supplementary Figure S3

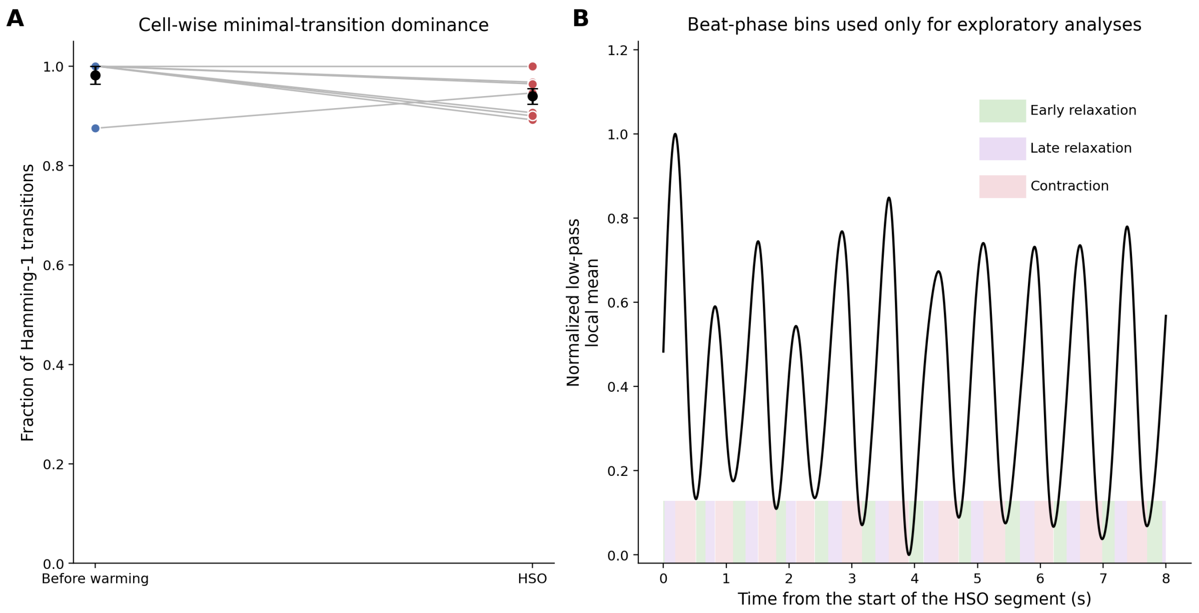


**Cell-wise support for the primary Hamming-1 result and beat-phase partition used only in exploratory analyses.** **A**, fraction of Hamming-1 transitions in each cell before warming and during HSOs. Every cell retained a high Hamming-1 fraction, showing that the main transition-topology result was not created by pooling across cells. This panel directly supports the topology-centered main claim. **B**, contraction, early-relaxation, and late-relaxation bins defined from the low-pass local 5-sarcomere mean trace in the exemplar HSO segment. These bins were used only for exploratory phase-bias tests; because they did not yield persuasive support in the present dataset, they were not advanced to the main text.

### Supplementary Figure S4

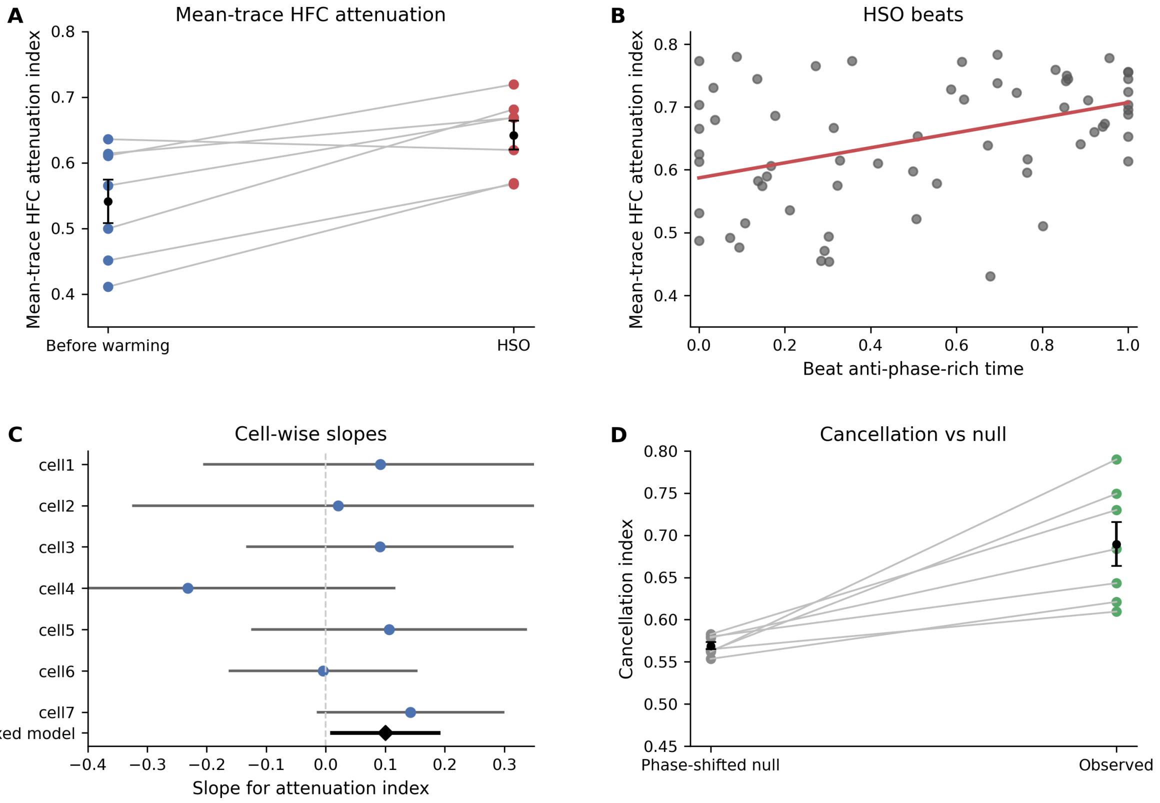


**Exploratory attenuation/cancellation analysis retained as supportive material. A**, paired values for the local 5-sarcomere mean-trace high-frequency attenuation index before warming and during HSOs. **B**, beat-level relation between anti-phase-rich time and attenuation index during HSOs; red line, fitted mixed-model fixed effect. **C**, cell-wise slopes for the beat-level attenuation model with the mixed-model estimate shown below. **D**, cell-wise comparison between the observed phase-vector cancellation index and a within-beat phase-shifted null during HSOs. In the revised manuscript hierarchy, this figure is retained to show supportive consistency with cancellation-like averaging, but it is not treated as a co-equal bridge result because the cycle-level $A \times R_w$ summary provides a cleaner and more transparent within-segment description of the mean fast signal.

### Supplementary Figure S5

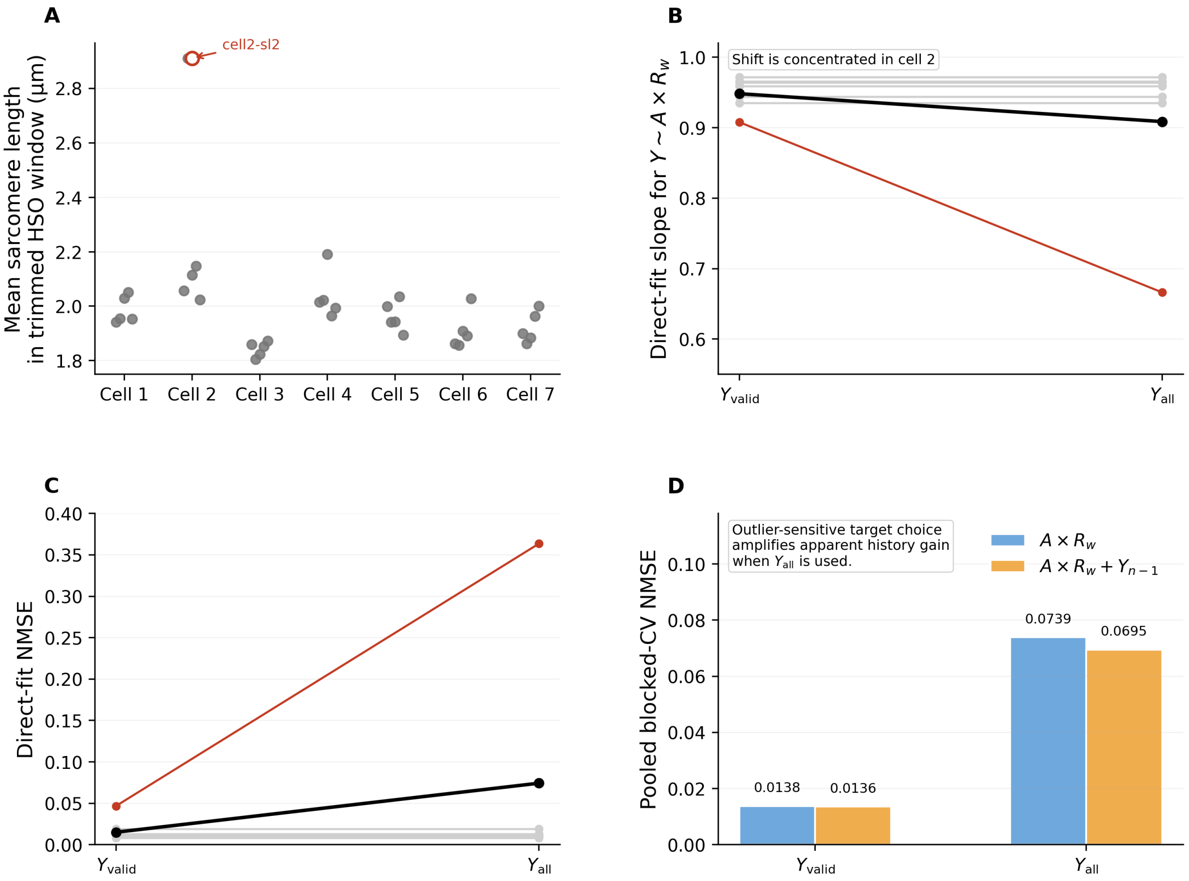


**Target-choice sensitivity for the descriptive cycle-level amplitude-synchrony bridge. A**, mean sarcomere length within the trimmed HSO window for every sarcomere trace used in the source dataset. Cell 2 sarcomere 2 (cell2–sl2) is the only trace clearly separated from the others in trimmed-window mean sarcomere length. **B**, direct-fit slopes for $Y \sim A \times R_w$ when the target is defined from the valid-sarcomere mean trace ($Y_{\mathrm{valid}}$) or from the all-sarcomere mean trace ($Y_{\mathrm{all}}$). The change is concentrated in cell 2, whereas the other cells remain nearly unchanged. **C**, direct-fit normalized mean squared error for the same comparison. Errors remain low for $Y_{\mathrm{valid}}$ but are inflated for $Y_{\mathrm{all}}$ because that separated trace enters the averaged target. **D**, pooled blocked-cross-validation normalized mean squared error for $A \times R_w$ and $A \times R_w + Y_{n-1}$ under the two target choices. For $Y_{\mathrm{valid}}$, the apparent gain from adding $Y_{n-1}$ remains tiny (0.0138 to 0.0136), whereas for $Y_{\mathrm{all}}$ it becomes larger (0.0739 to 0.0695) because the target is no longer aligned with the quality-control-valid sarcomere set used to define the explanatory variables. This figure therefore supports the descriptive-bridge interpretation and argues against stronger over-reading of history terms.

#### Supplementary Table S1. Paired robustness of the primary topology pillars and supportive attenuation summary

| Analysis set | Pre valid | HSO valid | Δ valid | $P$ | Δ anti-phase-rich | Δ attenuation |
|---|---|---|---|---|---|---|
| Fixed primary gate | 0.298 | 0.956 | 0.658 | 0.0156 | 0.255 (0.0156) | 0.101 (0.0312) |
| Refined display gate | 0.308 | 0.994 | 0.686 | 0.0156 | 0.259 (0.0156) | 0.101 (0.0312) |
| Refined display gate, excluding cell 2 | 0.301 | 0.993 | 0.692 | 0.0312 | 0.284 (0.0312) | 0.120 (0.0312) |

The increase in valid phase-trackable time and the increase in time with three or more anti-phase neighboring pairs are the primary paired robustness results carried into the main manuscript. The increase in the mean-trace high-frequency attenuation index is shown alongside them as supportive material. Values in parentheses are paired Wilcoxon $P$ values for the anti-phase-rich and attenuation effects.

#### Supplementary Table S2. Parameter robustness of the primary topology result and supportive attenuation summary

| Summary quantity across 54 settings | Value |
|---|---|
| HSO fraction of Hamming-1 transitions (range) | 0.876 to 0.980 |
| Secondary association coefficient for attenuation index (range) | 0.0746 to 0.1132 |
| Settings with positive secondary coefficient | 54/54 |
| Settings with $P < 0.05$ for the secondary coefficient | 53/54 |
| HSO valid time fraction (range) | 0.897 to 1.000 |

Sensitivity analysis over gate threshold, gap-fill duration, dwell time, and phase-smoothing window. The primary conclusion—dominance of Hamming-1 transitions—remained stable across the full parameter grid and therefore supports the topology-centered main claim. The attenuation association was also directionally stable, but remains supportive rather than primary in the revised hierarchy.

#### Supplementary Table S3. Supportive robustness of the attenuation association

| Model / predictor | $\beta$ | 95% CI | $P$ | Beats |
|---|---|---|---|---|
| Fixed primary gate, three-or-more anti-phase neighboring pairs | 0.099 | 0.020 to 0.179 | 0.0145 | 68 |
| Fixed primary gate, excluding cell 2 | 0.095 | 0.019 to 0.170 | 0.0142 | 58 |
| Refined display gate, three-or-more anti-phase neighboring pairs | 0.108 | 0.027 to 0.189 | 0.0088 | 68 |
| Refined display gate, excluding cell 2 | 0.096 | 0.018 to 0.173 | 0.0151 | 58 |
| Alternative definition: two or more anti-phase neighboring pairs | 0.056 | −0.052 to 0.163 | 0.3087 | 68 |
| Alternative definition: exactly three anti-phase neighboring pairs | 0.096 | −0.008 to 0.200 | 0.0714 | 68 |
| Alternative definition: four anti-phase neighboring pairs | 0.146 | 0.013 to 0.279 | 0.0317 | 68 |
| Alternative definition: mean anti-phase neighboring-pair count | 0.048 | 0.007 to 0.088 | 0.0220 | 68 |

The main anti-phase-rich definition used in the manuscript gave a stable positive association across gate choices and after exclusion of cell 2. Alternative definitions were more variable, which is why the attenuation/cancellation analysis is framed as supportive rather than as a uniquely determined mechanistic result or a main-text pillar.

#### Supplementary Table S4. Exploratory analyses not advanced beyond supportive status

| Analysis | Result summary | Interpretation |
|---|---|---|
| Beat-label shuffle null for the refined-gate secondary association | Observed $\beta = 0.108$; empirical upper-tail $P = 0.0357$ | Supports the secondary association; retained in Supplementary Figure S2. |
| Circular-shift null for the refined-gate secondary association | Observed $\beta = 0.108$; empirical upper-tail $P = 0.0288$ | Supports the secondary association; retained in Supplementary Figure S2. |
| Beat-phase dependence of Hamming-1 transitions | Friedman $P = 1.000$ across contraction, early relaxation, and late relaxation | No persuasive phase bias; not advanced to the main text. |
| Beat-phase dependence of anti-phase-rich occupancy | Friedman $P = 0.276$ across contraction, early relaxation, and late relaxation | Descriptively suggestive but not statistically supported; not advanced to the main text. |
| Beatwise frequency dispersion across the five sarcomeres | Mean dominant frequency 6.12 Hz; mean within-beat SD 0.259 Hz; mean CV 0.0439 | Useful descriptively, but did not yield a sufficiently clean complementary result; not advanced to the main text. |

Only analyses supported by the data were advanced to the main manuscript. Weakly supported or unsupported exploratory analyses, or analyses retained only as supportive material, are listed here explicitly so that the scope and ranking of the present evidence remain clear.

#### Supplementary Table S5. Expanded descriptive-bridge statistics for the canonical target $Y_{\text{valid}}$

| Cell | Cycles | $\beta_1$ | $r$ | Direct NMSE | Additive CV | $A \times R_w$ CV | $+ Y_{n-1}$ CV | $+ (A \times R_w)_{n-1}$ CV | + 2-step CV |
|---|---|---|---|---|---|---|---|---|---|
| cell1 | 45 | 0.972 | 0.990 | 0.0192 | 0.1352 | 0.0157 | 0.0156 | 0.0155 | 0.0158 |
| cell2 | 39 | 0.908 | 0.976 | 0.0468 | 0.1606 | 0.0596 | 0.0673 | 0.0691 | 0.0756 |
| cell3 | 49 | 0.959 | 0.994 | 0.0118 | 0.1065 | 0.0088 | 0.0101 | 0.0101 | 0.0099 |
| cell4 | 41 | 0.944 | 0.995 | 0.0104 | 0.1096 | 0.0173 | 0.0168 | 0.0170 | 0.0180 |
| cell5 | 44 | 0.935 | 0.994 | 0.0116 | 0.0655 | 0.0159 | 0.0145 | 0.0143 | 0.0148 |
| cell6 | 47 | 0.966 | 0.993 | 0.0131 | 0.1213 | 0.0181 | 0.0186 | 0.0186 | 0.0188 |
| cell7 | 43 | 0.964 | 0.996 | 0.0080 | 0.1144 | 0.0093 | 0.0103 | 0.0102 | 0.0115 |
| Pooled | 308 | 0.948 | 0.992 | 0.0151 | 0.1006 | 0.0138 | 0.0136 | 0.0135 | 0.0137 |

Cell-wise and pooled statistics for the canonical cycle-level target $Y_{\text{valid}}$. Direct fits summarize the descriptive benchmark $Y_{\text{valid}} \sim \beta_0 + \beta_1(A \times R_w)$, where $\beta_1$ close to 1 indicates near-proportional scaling within the same observed segment rather than proof of an independent law. Blocked-cross-validation normalized mean squared error (CV NMSE; smaller is better) is shown for the additive current-state model, the $A \times R_w$ summary, and the three history-augmented variants requested in review-oriented sensitivity checks: $A \times R_w + Y_{n-1}$, $A \times R_w + (A \times R_w)_{n-1}$, and $A \times R_w$ plus a two-step history summary. The pooled row shows that the main gain comes from moving from the additive model to the multiplicative amplitude-synchrony summary, whereas the history-augmented variants change error only marginally. This table is therefore intended as documentation of descriptive parsimony for the bridge analysis, not as evidence for a universal governing relation.
